# TDP-43 loss of function drives aberrant splicing in Parkinson’s disease

**DOI:** 10.1101/2025.09.04.673943

**Authors:** Jonathan W. Brenton, Jordan Follett, Raja Nirujogi, Christina E. Toomey, Patrica Lopez-Garcia, James R. Evans, Yeon J. Lee, Khaja Mohieddin Syed, Guillermo Rocamora Perez, Áine Fairbrother-Browne, Karishma D’Sa, Melissa Grant-Peters, Joanne Lachica, Amy R. Hicks, Aaron Z. Wagen, Benjamin O’Callaghan, Hannah Macpherson, Kylie-ann Montgomery, Oriol Busquets, Regina H. Reynolds, Sonia Garcia Ruiz, Tianyu Cao, Zhongbo Chen, Hélène Plun-Favreau, Philip C. Wong, Matthew Farrer, Tammaryn Lashley, Frank Soldner, Dirk Hockemeyer, Dario Alessi, Nicholas W. Wood, John Hardy, Donald C. Rio, Zane Jaunmuktane, Emil K. Gustavsson, Sonia Gandhi, Mina Ryten

## Abstract

While mRNA splicing dysregulation is a well-established contributor to neurodegeneration in disorders such as amyotrophic lateral sclerosis (ALS) and frontotemporal dementia (FTD), its role in Parkinson’s disease (PD) remains underexplored. Here, we analyse transcriptomic data from >500 post-mortem human brain samples from individuals with and without PD to show that splicing alterations are frequently detected. Differentially spliced genes were significantly more enriched for those causally-implicated in both PD and ALS than genes that were differentially expressed. Furthermore, we observed a strong association between these splicing alterations and dysfunction of the RNA-binding protein (RBP), TAR DNA-binding protein 43 (TDP-43). Strikingly, genes and exon junctions affected by TDP-43 knockdown overlapped significantly with those dysregulated across brain regions in PD. In brains from individuals with the *LRRK2* c.6055G>A (p.G2019S) mutation, the most common genetic cause of PD, we also observed significant enrichment of TDP-43-dependent splicing changes. This finding was corroborated in human pluripotent stem cell-derived midbrain dopaminergic neurons and a *LRRK2* p.G2019S knock-in mouse model, where reduced nuclear TDP-43 levels evidenced the well-recognised loss-of-function mechanism contributing to splicing dysregulation. By leveraging our RNA-based analyses we predicted TDP-43-dependent novel peptide sequences and validated their existence within human *LRRK2* mutation mDNs, while also demonstrating an overall loss of protein and mRNA expression in mis-spliced genes. Collectively, our findings reveal that PD is marked by extensive splicing dysregulation dependent on TDP-43, making TDP-43 a promising new therapeutic target in PD.

## Main

Alternative splicing has the potential to drive genome-wide alterations in transcript use and protein isoforms. To study its role in PD, we generated high-depth RNA-sequencing (RNA-seq) data designed to span multiple exons from post-mortem brain samples of individuals diagnosed with PD and clinically unaffected controls (∼110 million, 150 base pair (bp) unique genome-mapping reads per sample). We analysed two pathological stages of disease, mid- (Braak stage III and IV) and late-stage disease (Braak stage VI), across multiple brain areas, including the substantia nigra, basal ganglia and cortices (Supplementary Table 1). This unique dataset enabled us to comprehensively profile differential gene expression and alternative splicing across disease stages and brain areas (Fig. 1a). Strikingly, only a small proportion of genes (5.5%; Fig. 1b) were both differentially expressed and differentially spliced. Analysis of the significant gene sets from each method revealed a comparably low proportion of shared pathways (1.3%; Extended Data Fig. 1). This suggests that in idiopathic PD gene expression and splicing predominantly affect distinct biological processes.

**Fig. 1:**
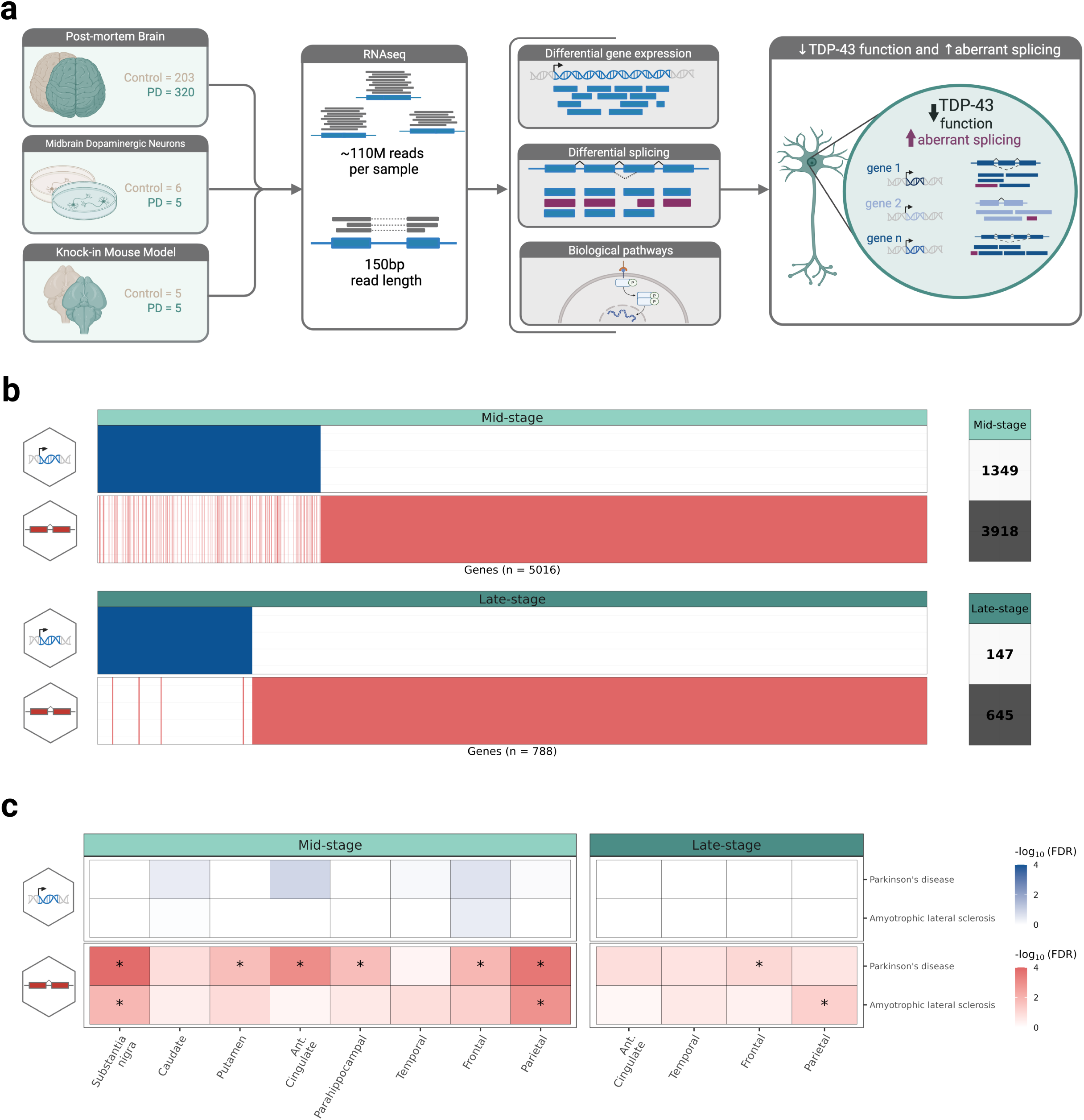
Splicing and gene expression in sporadic Parkinson’s disease exhibit distinct profiles, with splicing showing stronger disease associations. **a,** Overview of the study across different experiments. RNA-seq was conducted on multiple data sources, human post-mortem tissue (sporadic (Mid Stage: Braak stage 3-4; Late Stage: Braak stage 6) and *LRRK2* p.G2OI 9S PD cases), *LRRK2* p.G2OI 9S knock-in (KI) mouse and *LRRK2* p.G2OI 9S human pluripotent stem cell (hPSC) models to generate deeply sequenced and exon junction spanning reads that were used for differential gene expression (DE), differential splicing (DS) and gene ontology (GO) pathway analysis. Differentially spliced data was compared to a database of genes and junctions altered upon TDP-43 knockdown (KD) to demonstrate a loss of TDP-43 function, a known cause of neurodegeneration, in our cohorts, b, Binary plot of common genes found through DE and DS analyses in sporadic PD (FDR < 0.05). Each bar represents a single unique gene and rows indicate differentially expression (blue) or splicing (red) at different pathological stages of PD (panels). Total numbers of unique genes found for each stage and analysis shown to the right. Mid Stage: n=234 (Control n=113, PD=121), Late Stage: n=272 (Control n=8O, PD=192). **c,** Enrichment analysis of genes with a genetic association with PD and amyotrophic lateral sclerosis (ALS) in sporadic across brain regions and disease stage (* FDR < 0.05).

To evaluate the disease relevance of these findings, we investigated whether genes causally implicated in PD, identified through genome-wide association studies and Mendelian disease loci^1^, were enriched among the differentially expressed or spliced genes. Indeed, we observed a consistent enrichment of PD-associated genes among the differentially spliced genes, whereas no such enrichment was seen among the differentially expressed genes (Fig. 1c). Given that ALS is a neurodegenerative disease with a clear genetic association with splicing dysregulation due to disease-causative mutations in RBPs^2^, we also tested for the enrichment of genes causally implicated in ALS. Our analysis revealed a significant overrepresentation of ALS-associated genes among the alternatively spliced genes in PD, suggesting shared mechanisms of splicing dysregulation between the two disorders.

Since the loss of TAR DNA-binding protein 43 (TDP-43) splicing regulation is a central feature of ALS, we investigated whether this phenomenon also occurs in idiopathic PD. To define a TDP-43 loss-of-function signature, we generated a splicing database of junctions and genes alternatively spliced upon nuclear absence or knockdown (KD) of TDP-43 from eight independently generated experiments^3–8^ (Fig. 2a). We found a significant enrichment of these TDP-43 loss-of-function events among differentially spliced junctions and genes in idiopathic PD (Fig. 2b). Gene ontology (GO) pathway analysis of the differentially spliced genes shared between PD and TDP-43 loss-of-function indicates that synaptic and cytoskeletal processes are major targets of mis-splicing (Fig. 2c). Analysis of pathways encoding genetic risk for PD^9^ also highlighted endolysosomal function (vesicle-mediated transport and clathrin-mediated endocytosis pathways) as a further target of mis-splicing among other pathways related to neuronal structure and function (Fig. 2d). Since TDP-43 loss of function disrupts cytoskeletal and synaptic processes in ALS^3,10,11^, and that endolysosomal dysfunction is a core feature of PD, our findings suggest that TDP-43–dependent mis-splicing affects functionally important and key disease-relevant pathways in PD.

**Fig 2:**
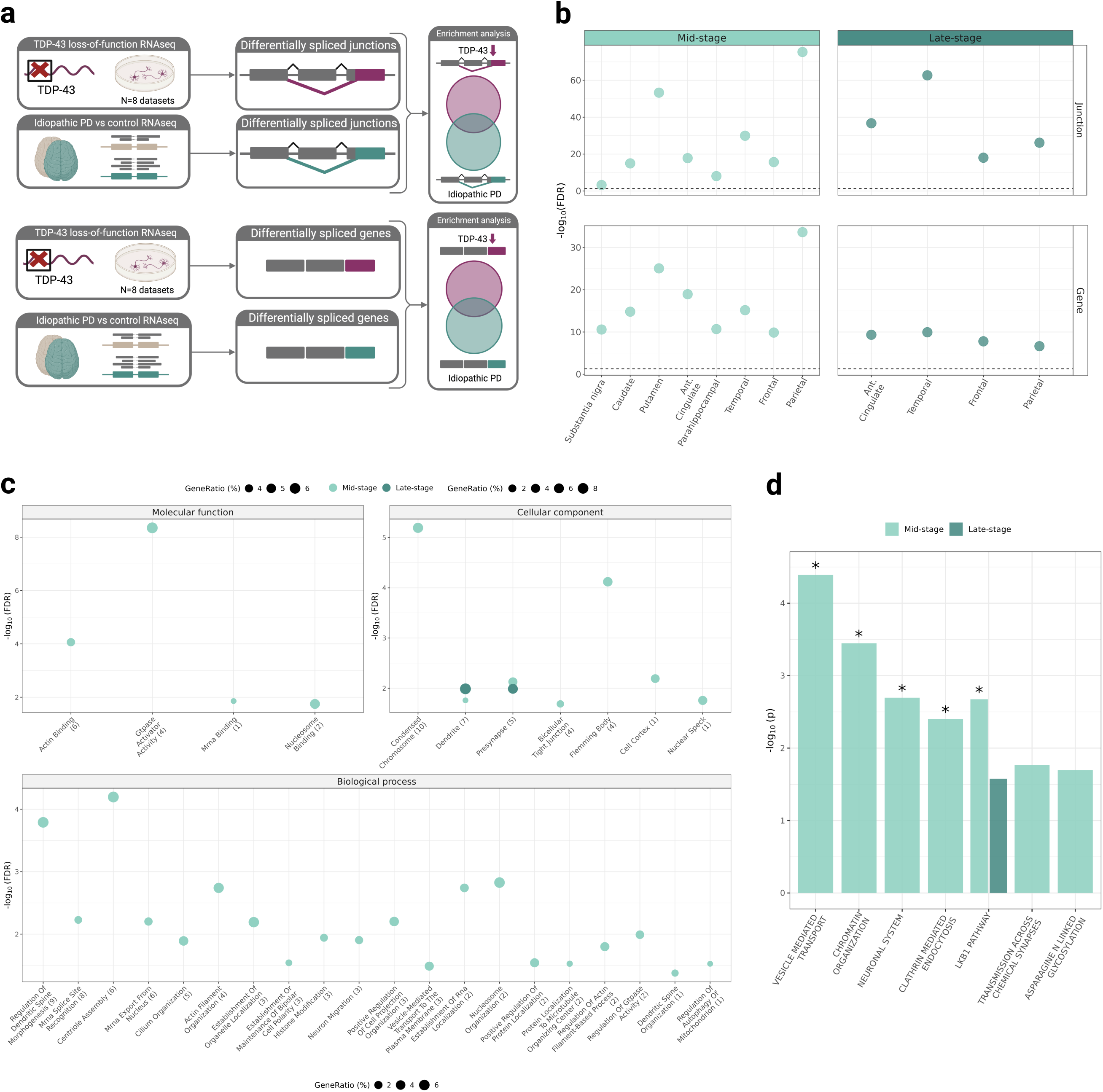
Nuclear TDP-43 loss of function in mid- and late-stage sporadic PD results in mis-splicing of synaptic genes. **a,** Diagram to illustrate the design of experiments assessing TDP-43-dependent splicing events and associated genes in sporadic PD. Differentially spliced junctions and genes found in post-mortem mid- and late-stage sporadic PD were compared to those observed in human TDP-43 knockdown (KD) models (termed TDP-43-dependent splicing events and genes), b, Dot plots to show the enrichment of TDP-43-dependent junctions (top row) and genes (bottom row) by stage and brain area (right panel, hypergeometric test, dotted line at FDR<0.05; Mid Stage: n=234 (Control n=113, PD=121), Late Stage: n=272 (Control n=8O, PD=192)). **c,** Dot plot to show significant GO pathway enrichments (FDR<0.05) amongst genes that were both differentially-spliced in human TDP-43 KD models and in mid- (light green) and late-stage PD (dark green). Semantic similarity was used to reduce the numbers of pathways displayed with the number of reduced terms indicated within parentheses, **d,** Bar plot showing enrichment in pathways encoding genetic risk for PD^9^. All nominal p-value significant pathways shown, with those passing FDR-adjustment indicated with an asterisk.

Next, we investigated whether TDP-43 dysfunction contributes to pathogenesis in PD cases harbouring the LRRK2 p.G2019S mutation. We focused on this form of PD, not only because it the most common type of Mendelian PD, but because of reports of increased TDP-43 pathology in LRRK2 PD^12,13^, and recent studies demonstrating a significantly higher proportion of a-synuclein seeding amplification (SAA) negative CSF samples among individuals with LRRK2 pG2019S-positive PD compared to sporadic cases^14,15^. We performed high-depth RNA-sequencing on post-mortem brain tissue from two regions in patients with *LRRK2* p.G2019S-associated PD to assess TDP-43–dependent mis-splicing (Fig. 3a). This revealed strong evidence of TDP-43 loss-of-function, as TDP-43-sensitive junctions and genes were significantly enriched among those identified through differential splicing analyses in the *LRRK2* p.G2019S PD brain samples (Fig. 3b, left panels). This finding was further investigated in two separate RNA-seq datasets generated from human pluripotent stem cell (hPSC)-derived midbrain dopaminergic neurons (mDNs) with the LRRK2 p.G2019S mutation (Fig. 3a). These results confirmed that the *LRRK2* p.G2019S mutation led to an increase in TDP-43-dependent mis-splicing in these cells (Fig. 3b, right panels). GO and genetically-implicated pathway analyses of *LRRK2* p.G2019S-positive brains and cell models revealed biological processes similar to those observed in idiopathic PD. They revealed TDP-43-dependent splicing dysfunction in synaptic and cytoskeletal genes (Fig. 3C), as well as in genes involved in endolysosomal function (vesicle-mediated transport, clathrin-mediated endocytosis and lysosome pathways; Fig. 3D). We also examined total and phosphorylated TDP-43 protein levels and found no significant increases in the frontal cortex of LRRK2 p.G2019S or idiopathic PD cases (Extended Data Fig. 2; Supplementary Table 2). This is consistent with previous studies showing that TDP-43-dependent splicing dysregulation can occur in the absence of detectable TDP-43 aggregates^16,17^.

**Fig. 3:**
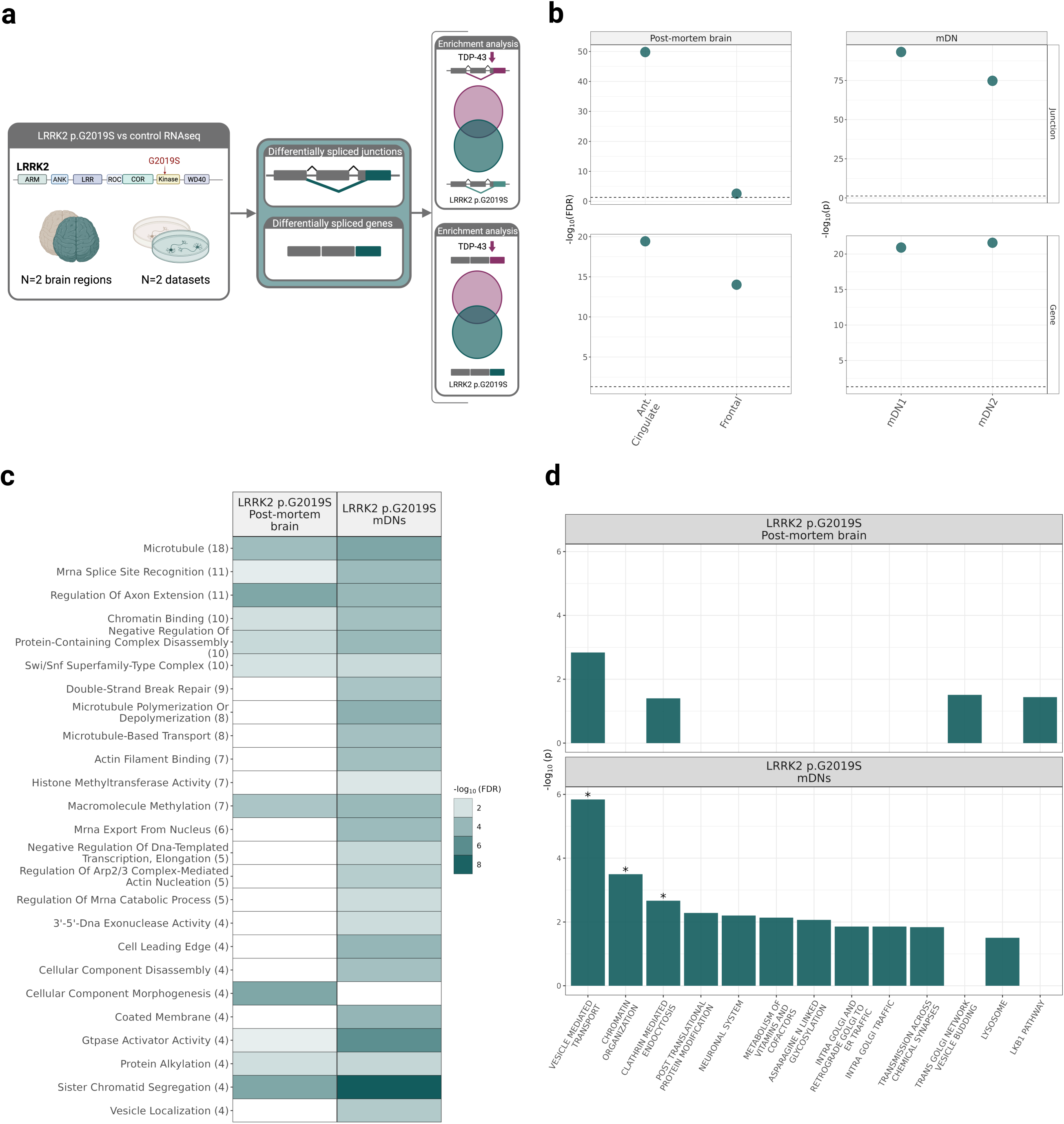
Nuclear TDP-43 loss of function and associated mis-splicing is a feature of PD in *LRRK2* p.G2OI9S mutation carriers. **a,** Overview of *LRRK2* p.G2OI 9S experiments, **b,** Dot plots to show the enrichment of TDP-43 dependent splicing events and genes amongst those detected in both post-mortem human brain tissue from individuals with PD due to the *LRRK2* p.G2OI 9S mutation (n=17 (Control=l 0, *LRRK2* p.G2OI 9S = 7)), and in hPSC-derived midbrain dopaminergic neurons generated from patients carrying the same mutation (dotted line denotes p = 0.05; mDN1: n=6 lines, mDN2: n=5 lines). Significant enrichments were found in all post-mortem human brain regions tested and in two independently generated midbrain dopaminergic neuron models, c, Heatmap to show top 25 central GO pathway enrichments (FDR <O.O5) amongst genes that were both differentially-spliced in human TDP-43 knockdown models and in *LRRK2* p.G2OI 9S post-mortem cases. Semantic similarity was used to reduce the numbers of pathways displayed with the number of reduced terms indicated within parentheses, **d,** Bar plot showing enrichment in pathways encoding genetic risk for PD^9^. All nominal p-value significant pathways shown, with those passing FDR-adjustment indicated with an asterisk.

To study the impact of the *LRRK2* p.G2019S mutation in vivo, we investigated splicing activity in the well-characterized *LRRK2* p.G2019S knock-in (KI) mouse model ^18–21^. Given the notable species differences in TDP-43 splicing between humans and mice^22^, we leveraged five publicly available TDP-43 KD datasets to compile a mouse-specific database of TDP-43-dependent junctions and genes^23–27^ (Fig. 4a). Using this database, we observed significant enrichment of TDP-43-sensitive loss-of-function events among the junctions and genes that were differentially spliced in the *LRRK2* p.G2019S KI mouse brain (Fig. 4b). We also investigated TDP-43 protein distribution in primary cortical neurons from *LRRK2* p.G2019S KI mice by confocal microscopy (Fig. 4c). This revealed a significant reduction in nuclear TDP-43, whereas the distribution of the RBP FUS remained similar to control neurons (Fig. 4d). In parallel, western blot analysis of subcellular fractions confirmed the selective decrease in nuclear TDP-43 protein in these *LRRK2* p.G2019S primary neurons (Fig. 4e). These results specifically demonstrate the well-established loss-of-function mechanism through which TDP-43 nuclear loss drives mis-splicing, providing a mechanistic basis for the splicing abnormalities detected in *LRRK2* p.G2019S models and in PD.

**Fig. 4:**
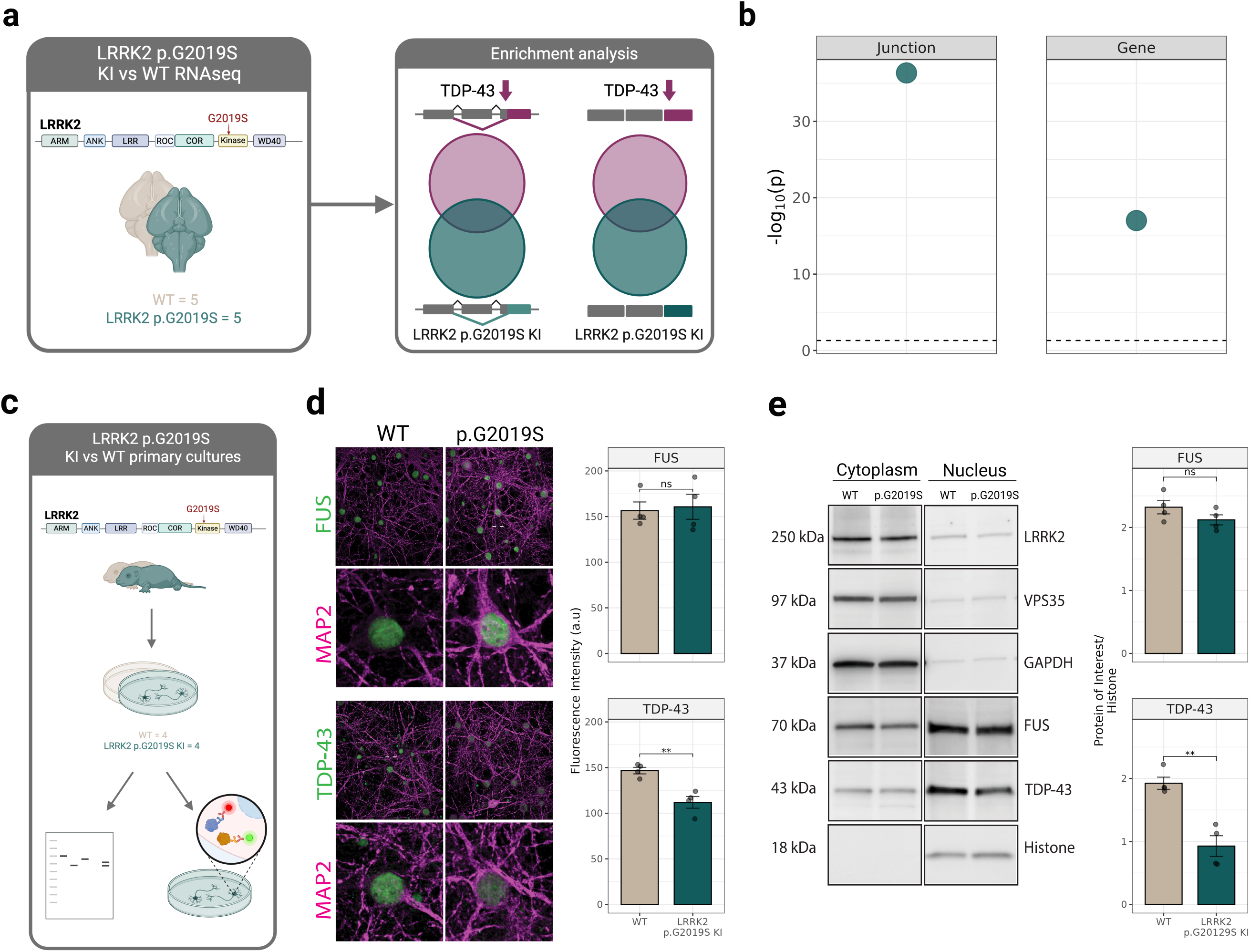
The *LRRK2* p.G2OI 9S KI mouse model exhibits TDP-43 dependent mis-splicing with concurrent loss of nuclear TDP-43 protein in neurons. **a,** Overview of the RNA-sequencing analyses conducted in the *LRRK2* p.G2OI 9S KI mouse model. Brains from 3-6 month old animals (n=10 (WT=5, *LRRK2* p.G2OI 9S = 5)) were analysed for the enrichment of TDP-43 dependent splicing events and associated genes, b, Dot plots to show the enrichment of TDP-43 dependent junctions and linked genes (dotted line at p = 0.05). **c,** Overview of analyses of nuclear TDP-43 protein in the *LRRK2* p.G2OI 9S KI mouse model. Primary cultures of cortical neurons from P0 pups were analysed using western blotting and immunocytochemistry to assess TPD-43 nuclear protein levels (n=8 separate cultures (WT=4, *LRRK2* p.G2OI 9S = 4)). d, Immunocytochemical analysis of TDP-43 nuclear protein levels in cortical neurons. Representative images (left) from primary cultures stained for FUS (top panel) or TDP-43 (bottom panel). Quantification of nuclear TDP-43 staining shown to the right, e, Western blotting for nuclear TDP-43 protein levels in cortical neurons. Representative blots (left) of protein levels following nuclear and cytoplasmic extractions. Quantification of nuclear FUS (top panel) and TDP-43 (bottom panel) shown to the right. ** = p<0.01, ns = p>0.05.

Next, we focused on the potential consequences of TDP-43-dependent cryptic splicing events, namely i) the generation of de novo proteins, and ii) reduced protein expression secondary to RNA instability and degradation. We focused on one *LRRK2* p.G2019S mDN dataset (mDN1) that showed clear evidence of TDP-43-dependent mis-splicing and included matched RNA-seq and mass spectrometry data (Fig. 5a). To detect evidence of cryptic peptides, defined here as peptides containing TDP-43-dependent mis-splicing junctions, we generated RNA sequences incorporating novel mis-splicing events and predicted alternative open reading frames (ORFs) encoding novel peptides. We focused on the detection of peptides with high mapping specificity to the novel ORF as compared to the known protein sequence. This approach revealed the significant increase in the translation of several predicted cryptic peptides in *LRRK2* p.G2019S mDNs (false discovery rate-adjusted p-value (FDR) < 0.05; Fig. 5b), providing evidence of aberrant protein expression due to TDP-43-dysfunction. We next assessed the impact of TDP-43-mediated mis-splicing on both differential gene and protein expression using the same paired datasets (Fig. 5c). Genes with TDP-43–dependent mis-splicing events were more likely to be downregulated at both the mRNA and protein levels (>50%; p < 0.01; Fig. 5d). Furthermore, previously identified mis-spliced genes^28^ were exclusively found in this subset of genes downregulated at both the transcript and protein levels (Extended Data Fig. 3). Our findings indicate that TDP-43–dependent mis-splicing in PD drives aberrant cryptic peptide expression and a widespread reduction in both transcript and protein abundance, with implications for cellular dysfunction.

**Fig. 5:**
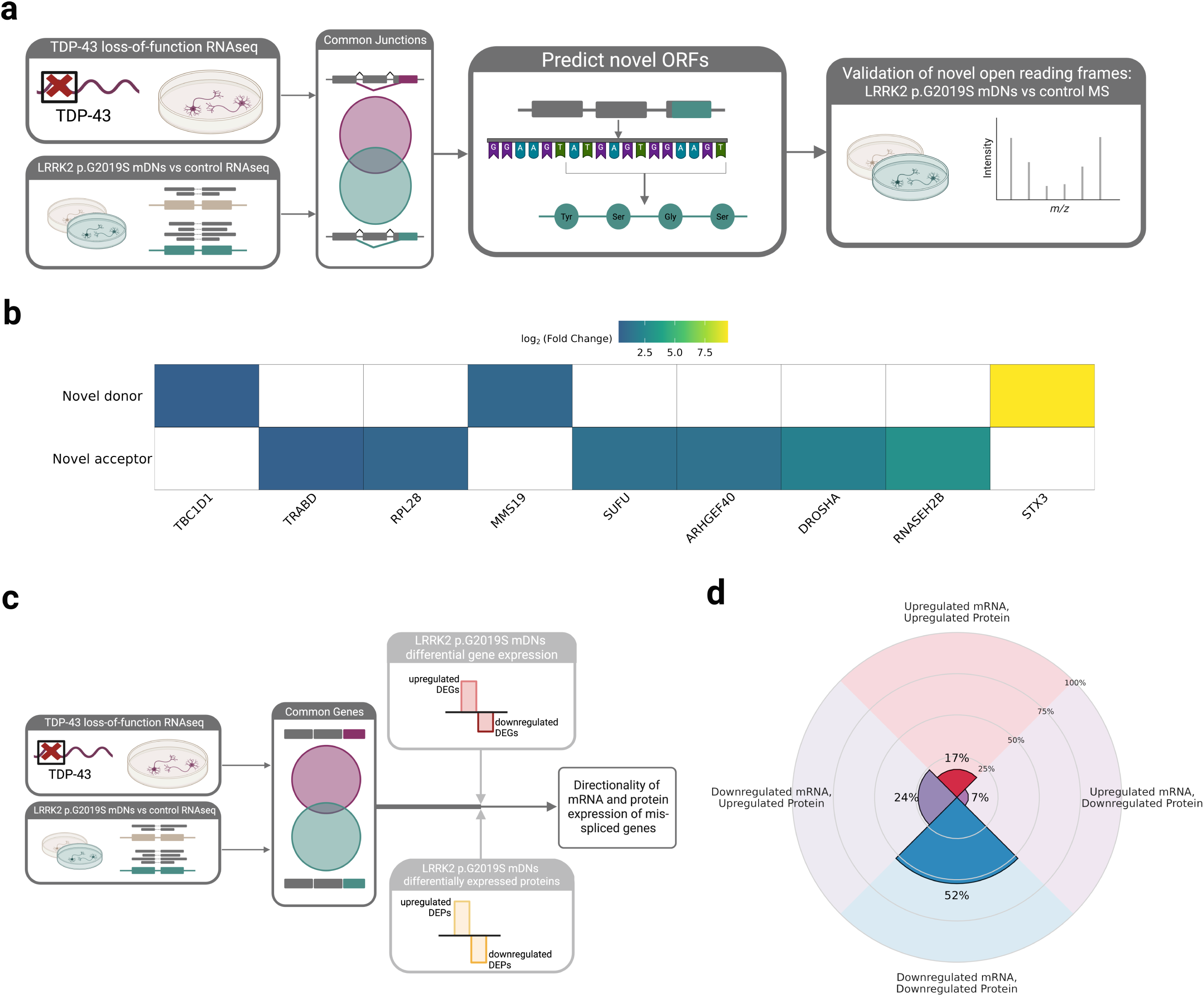
TDP-43 dependent cryptic splicing in *LRRK2* p.G2OI9S mDNs produces cryptic peptides and a downregulation of mRNA and protein in target genes. **a,** Overview of cryptic peptide detection analysis conducted in a hPSC LRRK2 p.G2OI9S mDN model (n=5 lines (Control=3, *LRRK2* p.G2OI9S=2, 6 replicates per line)). TDP-43-dependent splicing events were converted into open reading frames by replacing annotated junctions with novel donor or acceptor sites, and translated into peptide sequences. Paired mass spectrometry (MS) data was used to validate the presence of cryptic peptides, by aligning resulting peptide ions to the novel peptide sequences, **b,** Heatmap illustrating genes with upregulation in high confidence cryptic peptides (FDR < 0.05). **c,** Diagram of analyses of mRNA and protein levels in TDP-43 mis-spliced genes. To assess the impact and directionality of TDP-43-dependent mis-splicing, genes exhibiting TDP-43 mis-splicing and differential expression at both mRNA and protein levels were examined, **d,** Circular bar plot illustrating that TDP-43-dependent mis-splicing leads to a predominant downregulation of target genes at both an mRNA and protein level (p<0.05).

Together, our data establish TDP-43–dependent mis-splicing as a consistent and disease-associated feature of PD. TDP-43–dependent mis-splicing is found across multiple stages of idiopathic PD, as well as in *LRRK2* p.G2019S-positive disease, the most common monogenic form. Furthermore, this phenomenon is evident in multiple mDN models and a mouse model of *LRRK2*-positive PD. The latter is associated with a concomitant loss of nuclear TDP-43 protein, providing a mechanistic explanation for the findings. As would be expected, the mis-splicing events we observed in PD were accompanied by the expression of cryptic peptides, and loss of mRNA and protein levels. Furthermore, we note that the genes mis-spliced due to TDP-43 loss-of-function in idiopathic and *LRRK2* p.G2019S PD were primarily related to the synapse and the cytoskeleton, suggesting that neuronal activity and structure could be severely impacted, as in ALS^5,6,10^.

The identification of TDP-43-dependent mis-splicing in PD is particularly important given the increasing recognition that nuclear TDP-43 loss-of-function could contribute to a range of diseases. While TDP-43 is commonly associated with ALS-FTD, its aggregation and loss of function have also been frequently observed in other major neurodegenerative disorders, including Huntington’s disease, and Alzheimer’s Disease. In fact, in Alzheimer’s disease, TDP-43 loss-of-function activity was demonstrated even in the absence of detectable protein aggregation^16,17^, a pattern we now show also occurs in PD. This indicates that mis-splicing can occur prior to or potentially independently of cytoplasmic aggregation of TDP-43. Thus, our findings contribute to an important and growing literature suggesting a wider role for TDP-43 dysfunction across neurodegenerative diseases that includes movement disorders.

In this context, our findings relating to the role of TDP-43-dependent mis-splicing in *LRRK2*-associated forms of PD are particularly significant. Despite the small sample numbers available for *LRRK2*-positive cases, we observed a highly significant enrichment of TDP-43-dependent mis-splicing events in brain tissue. This was corroborated by human cell models and the *LRRK2* p.G2019S KI model of disease. Given that it is well recognised that a sub-population of *LRRK2* p.G2019S PD patients do not show α-synuclein aggregation in their brains post-mortem^13^ and are negative on seed amplification assays^14,15^, our findings could suggest that TDP-43 loss-of-function could be a primary driver of disease in these cases. This could also explain the pleomorphic pathology observed in this subgroup, warranting further investigation.

This study has important implications for both the diagnosis and treatment of PD. The predictable detection of cryptic peptides in patient brain samples and cell models raises the question of whether it is possible to identify subsets of PD patients for whom TDP-43-dependent dysfunction is a major contributing factor. As has been demonstrated in the field of ALS-FTD, developing this type of novel set of diagnostic biomarkers will require additional research given that cryptic exon expression can be highly variable across brain regions, and these peptides are often present at low levels^29^. This presents a significant challenge when looking for these biomarkers in other tissues such as blood or CSF. Nonetheless, there is considerable interest in the identification of translated proteins containing cryptic exons as biomarkers for disease suggesting that these concerns are surmountable^30–33^.

Similarly, additional analyses are required to determine the value of correcting TDP-43-dependent mis-splicing in the context of PD. Recent work on ALS-FTD has found that loss of expression of key synaptic proteins through cryptic exon use can induce synaptic and axonal pathology and that correcting mis-splicing reverses these deficits^5,10^. Our analyses consistently revealed enrichment of synaptic and cytoskeletal-related function among the genes affected by TDP-43-dependent mis-splicing in PD, suggesting a shared mechanism across these neurodegenerative diseases that may be amenable to a common therapeutic strategy. In summary, our findings reveal a novel role for TDP-43 in PD pathogenesis and identify a potential therapeutic target and direction for future research.

## Methods

### Post-Mortem Brain Tissue Acquisition and Cohort Selection

PD cases were selected based upon neuropathological assessment and clinical diagnosis found in the clinical notes in patient records. All cases were assessed for α-synuclein pathology and rated according to Braak staging^34^. Idiopathic PD cases were categorized as mid-stage (Braak stage 3 and 4) or late-stage (Braak stage 6). Samples for the study of mid-stage PD were provided by the Parkinson’s Disease UK Tissue Bank (Imperial College London). Eight brain regions (substantia nigra, caudate, putamen, anterior cingulate cortex, temporal cortex, parahippocampal gyrus, frontal cortex and parietal cortex) were selected to provide a measure of disease progression from pathologically highly affected subcortical to unaffected cortical areas. Samples for the study of late-stage PD were sourced from the Queen Square Brain Bank for Neurological Disorders (UCL Queen Square Institute of Neurology). In this case, samples were selected from four cortical areas (anterior cingulate cortex, temporal cortex, frontal cortex and parietal cortex), providing a range of pathological aggregate load across regions.

Samples for the study of PD caused by the inheritance of the pathogenic *LRRK2* variant, p.G2019S, were provided by the Queen Square Brain Bank for Neurological Disorders (UCL Queen Square Institute of Neurology), and consisted of anterior cingulate and frontal cortical areas common to both idiopathic cohorts. The brain areas studied in each cohort, disease state and demographic details are provided in Extended Data Fig. 4 and Supplementary Table 1.

### Ethical Approval

Ethical approval for the study was obtained from the Local Research Ethics Committee of the National Hospital for Neurology and Neurosurgery (23/LO/0044). Informed consent was given to use the postmortem brain material for research in all cases.

### Tissue Preservation for Transcriptomic Analysis

Brain samples used in RNAseq analyses were flash-frozen at the time of post-mortem collection and stored at -80°C until use. Regional sampling dissections were performed on dry ice^35^.

### Bulk RNA-seq Library Preparation from Human Brain Tissue

RNA extraction from human post-mortem samples was performed using approximately 30-50 mg of frozen cortical or 10-30 mg of subcortical tissue per brain sample. The RNeasy 96 Kit (Qiagen) was used with the on-membrane DNase treatment step, as per manufacturer instructions, for tissue lysis and RNA isolation. Samples were quantified by absorption on the QIAxpert (Qiagen), and RNA integrity number (RIN) measured using the Agilent 4200 TapeStation (Agilent Technologies).

Stranded cDNA libraries from mid-stage and late-stage idiopathic PD samples were constructed with the KAPA mRNA HyperPrep Kit (Roche). 500ng of total RNA was used as input for each sample. xGen Dual Index UMI Adapters (Integrated DNA Technologies, Inc.) were added to each sample to minimise read mis-assignment when performing post-sequencing sample de-multiplexing.

*LRRK2* p.G2019S cohort cDNA libraries were prepared using a proprietary kit (Novogene UK), which involves the following steps: mRNA purification using poly-T oligo-attached magnetic beads, fragmentation, reverse transcription, second strand cDNA synthesis containing dUTPs, size selection and PCR amplification. cDNA libraries were multiplexed on S2 or S4 NovaSeq flow cells for paired-end 150 bp sequencing on the NovaSeq 6000 Sequencing System (Illumina) to obtain a target read depth of 110 million paired-end reads per sample. Sequenced reads were de-multiplexed and FASTQ files were generated using the BCL Convert software (Illumina).

### Processing of Human Brain RNA-seq Data

#### Quality Control, Transcriptome and Genome Alignment

FASTQ files were run through a Nextflow^36^ pipeline that performed quality control (QC) of reads and aligned the samples to the human genome and transcriptome (GRCh38 and Gencode v41 annotations^37^). Briefly, initial QC was performed using fastp^38^ (v0.23.2), which removed low quality reads and bases, as well as the trimming of adapter sequences. Filtered reads were aligned to the transcriptome using Salmon (correcting for sequence-specific, fragment GC-content and positional biases; v1.9.0; ^39^ using the entire genome as a decoy sequence. For splicing analyses, reads were aligned to the genome using STAR (v2.7.10a, with two-pass mode). FastQC^40^ (v0.11.9) was used to generate read QC data before and after fastp filtering. RSEQC^41^ (v4.0.0), Qualimap^42^ (v2.2.2a) and Picard^43^ (v2.27.5) were used to generate alignment QC metrics. MultiQC^44^ (v1.13) was used to visualise and collate quality metrics from all pipeline modules. Full details for the pipelines and parameters used for each module can be found here^45^.

https://github.com/Jbrenton191/RNAseq_splicing_pipeline.

Samples with a low read count (<60 million reads aligning to the transcriptome for post-mortem cohorts or > 30 million for the mouse cohort), a high proportion of reads aligning to intronic regions of the genome (>30%) or a deviant average GC content (<40% or <65%) were removed from further analysis.

Transcriptome aligned reads generated from Salmon were used to identify covariates and then outlier samples. For both processes, salmon read files were imported into R (v4.3.0) and converted into gene-level count matrices using the tximport (v1.30.0) and DESeq2 (v1.42.1) packages.

#### Covariate and Outlier Detection

Sample and sequencing covariates were assessed for collinearity using a custom script to reduce the number of covariates tested. This employed the stats (v4.2.3) and Hmisc (v5.2-0) packages to generate Spearman correlations, Kruskal-Wallis and chi-square tests to assess relationships between covariates. Of the numeric covariates that were collinear, those with the lowest overall correlations to all other variables were kept. VariancePartition (v1.32.5) scoring and correlations between remaining covariates and the top 10 principal components (PCs) were then used to assess which covariates should be controlled for in the final analysis.

Outlier samples were identified using these dataset-dependent covariates. Covariate-corrected gene counts, corrected using the limma^46^ (v3.58.1) removeBatchEffect function, were used to identify any sample that had a zscore > 3 on any of the top PCs (selected based on a cumulative variance cut off) or a z-score > 2 on its sample connectivity value calculated using the WGCNA^47^ package (v1.73). Both scores were calculated relative to the z-scores for all samples from that respective brain region.

### Transcriptomic and Functional Analysis of Bulk RNAseq Data

#### Differential gene expression

Gene-level count matrices were filtered for genes with 0 counts in at least one sample in either the disease or control group, as well as any gene overlapping ENCODE blacklist regions of the genome^48^. Analysis was performed using limma/voom^49^. Briefly, voom was used to shrink count dispersion and the duplicateCorrelation function was used to remove the effect of each individual to allow multiple brain areas from the same individual to be modelled together. These functions were applied twice as recommended by the developers. Group and brain area factors were merged into a single term to allow the model to be run once and all comparisons to be extracted from that model. Differentially expressed genes were extracted for each contrast using the eBayes and topTable limma functions and were classified as differentially expressed if their Benjamini and Hochberg FDR adjusted p-value was ≤ 0.05.

#### Differential Splicing Analysis

LeafCutter^50^ (v0.2.9) was used to perform differential splicing. SJ.out files containing junction reads were first converted to junction files with junctions overlapping ENCODE blacklist regions removed using a custom R script. Junction files were then clustered using the leafcutter_cluster.py script from the LeafCutter authors. The following options were used to remove junctions/introns with the following criteria: junctions supported by < 30 reads across all samples, supported by < 0.1% of the total number of junction read counts for the entire cluster and larger than 1 Mb.

Pairwise group comparisons, correcting for identified covariates, were made using LeafCutter’s differential_splicing function with the following options: intron clusters with more than 40 junctions and a coverage of ≤ 20 reads in ≤ 3 samples in per group were removed, as well as individual introns that were not found in ≥ 5 samples. Clusters were considered significant if they had an FDR-corrected p value of ≤ 0.05.

### Pathway Enrichment Analyses

#### Gene Ontology

Gene Ontology^51^ (GO) pathway analysis was performed using clusterProfiler (v4.10.1). The compareCluster function was used to perform over-representation analyses for all ontologies across brain areas or stages of disease. Background lists or universes consisted of any gene that passed initial filtering and was tested in either differential expression or splicing analyses. Pathways were counted as significant if their FDR-adjusted p-value was ≤ 0.05.

The number of GO pathways were reduced by semantic similarity using the GOSemSim^52^ and rrvgo^53^ (v1.14.2) R packages. The “Wang” method, a graph-based measure of semantic similarity, and a similarity threshold for grouping terms of 0.7 were used.

#### Genetic Risk Association

Gene lists from 46 biological pathways identified as linked to PD risk through common genetic variation^9^ was obtained from

https://pdgenetics.shinyapps.io/pathwaysbrowser/.

This collection contained three Pathway Interaction Database terms, eight KEGG terms and 35 Reactome terms. Fisher’s exact tests (one-sided) were used to test for enrichment of significant gene lists and were FDR-corrected by the number of pathways tested.

### Identification of Human TDP-43-Sensitive Junctions and Genes

Publicly available FASTQ files from the following datasets, PRJEB42763^3^ (all 3 separate experiments), PRJNA311234^4^, PRJNA497804^5^, PRJNA503396^6^, PRJNA522295^7^, PRJNA562297^8^, which performed experiments knocking down TDP-43 or comparing TPD-43 positive and negative nuclei. Only control and TDP-43 knockdown samples were analysed from within these datasets. FASTQ files were downloaded from GEO or ENA using sra_tools, ncbi_tools and entrez-direct packages. Any technical replicate FASTQs were merged and run with the nextflow alignment pipeline and differential splicing was performed as described above. Unique differentially spliced junctions and genes were collated across experiments to form a TDP-43-sensitive junction database.

### Enrichment Analyses of Disease-Linked Genes and TDP-43-sensitive Junctions and Genes

#### Enrichment of OpenTargets Disease Gene Lists

To assess the enrichment of genes associated with PD and ALS, genes identified by GWAS and functional genomics curated by OpenTargets^1^ (v25.03) were used as target gene lists. Gene lists for “Parkinson’s disease” (MONDO_0005180) and “Amyotrophic lateral sclerosis” (MONDO_0004976) were downloaded from the OpenTargets platform (https://platform.opentargets.org) and filtered for genes with scores above 0.1 in any of the genetic association data sources, to test genes that had causal evidence linking them to each disease.

#### Enrichment of TDP-43-sensitive Junctions and Genes

The TDP-43-sensitive junction database was used to identify genes and junctions that exactly matched those in the PD LeafCutter analyses, with junction matches based on exact start, end, strand, and chromosome location.

#### Enrichment Statistics

Fisher’s exact tests (one-sided) were used to test for enrichment of the target genes or junctions. The overlap of target junctions or genes with differentially expressed or spliced genes or junctions was compared to that of a background list of any unique significant or non-significant gene or junction tested in all brain areas in that cohort.

#### Neuropathological Characterization of TDP-43

Neuropathological assessments were performed using 8 μm mounted Formalin-Fixed Paraffin-Embedded (FFPE) tissue sections. These were dried at 37°C and baked at 60°C overnight prior to immunohistochemistry. Immunohistochemistry for TDP-43 was performed using methods published previously^54,55^. Sections were deparaffinised in xylene and rehydrated in decreasing grades of alcohol. Slides were incubated in a methanol/hydrogen peroxide (0.3%) solution for 10 min to block endogenous peroxidase activity. Sections were incubated in 98% formic acid for 10 minutes, followed by heat-induced epitope retrieval using 0.1 M citrate buffer (pH 6.0) for an additional 10 minutes. To block non-specific binding, sections were incubated in 10% bovine serum albumin for 30 minutes at room temperature and pressure (RTP). Sections were incubated in primary antibody (TDP-43 Proteintech 10782-2-AP 1:1000) for 1 hour at RTP. This antibody binds to both physiological TDP-43 and post-translationally modified TDP-43, including phosphorylated TDP-43. Sections were washed 3 times for 5 minutes in Tris-buffered saline with Tween and then were incubated for 30 min at RTP in biotinylated goat anti-mouse IgG secondary antibody (Vector Laboratories BA 9200, 1:200). Slides were washed as in the previous step and then incubated in pre-conjugated streptavidin–biotin complex (ABC) for signal amplification. Slides were then washed and incubated in 3,3ʹ-Diaminobenzidine (DAB) chromogen for 5 min then counterstained in Mayer’s haematoxylin. Finally, slides were dehydrated in increasing grades of alcohol (70, 90 and 100%), cleared in xylene and mounted with DPX. Slides were digitally scanned using a VS120 Olympus slide scanner. Images were loaded into QuPath and assessed for the presence of TDP-43 aggregation. Representative images were taken from each case.

### Generation of Human hPSC-derived Dopaminergic Midbrain Neurons with the *LRRK2* p.G2019S Mutation

Midbrain dopaminergic neurons generated from different human-induced pluripotent stem cells (hiPSCs) and human embryonic stem cell (hESC) lines were independently differentiated into dopaminergic neurons using established protocols.

#### Dataset 1

A fully characterized and sequenced feeder-free female hESC line (WIBR3; NIH approval number NIHhESC-10-0079) was differentiated into dopaminergic midbrain neurons as previously described^56^.

Three clonal *LRRK2* p.G2019S mutation cell lines were generated through TALEN or prime editing-based genome engineering of WIBR3 hESCs as previously described^56,57^, along with three control lines that underwent the editing process but do not exhibit any genetic modifications at the targeted locus.

Full differentiation^58^ and nucleofection protocols^59^ can be found on protocols.io.

#### Dataset 2

hiPSC lines from control and *LRRK2* p.G2019S carriers (Supplementary Table 3) were used to generate midbrain dopaminergic neurons using a small molecule-based protocol, as described previously^60^.

### RNA Extraction and Generation of RNA-seq Data from Dopaminergic Neurons

#### Dataset 1

At day 35 post differentiation into mDNs, RNA was isolated using the RNAeasy Minikit (Qiagen, 74104), followed by 30 min. of DNase treatment (Ambion, AM2238) at 37°C, isolation of poly(A)+ RNA transcripts [NEBNext poly(A) mRNA magnetic isolation module; New England Biolabs, E7490] from 0.5 µg of total RNA for RNA library preparation and sequencing using NEBNext Ultra Directional RNA Library Preparation Kit for Illumina (New England Biolabs, E7420S) according to the manufacturer’s instruction. Samples were sequenced using an Illumina S4 flow cell with 150 bp paired-end reads at the Vincent J. Coates Genomics Sequencing Laboratory at the University of California at Berkeley. Typical samples had a read depth of ∼200-250 million.

#### Dataset 2

At 5 weeks post-differentiation RNA was harvested from snap-frozen cell pellets using the Maxwell® RSC simply RNA Cells kit (Promega), and the accompanying Maxwell® RSC 48 instrument. After RNA extraction, the RNA concentration and quality using the 260/280 ratio were assessed using the nanodrop.

Sequencing libraries were prepared with the NEBNext Ultra II Directional PolyA mRNA Prep kit (NEB) using at least 250 ng of RNA. Fragmentation and PCR steps were undertaken as per the manufacturer’s instructions. Libraries were sequenced using NovaSeq S2 (Illumina) 200 cycles to generate 100 bp paired-end reads with an average read depth of ∼174 million paired-end reads per sample.

### Processing and Analysis of RNAseq data from hPSC Dopaminergic Neurons

In both datasets, replicates from different differentiations were merged into one sample per line/donor.

#### Dataset 1

FASTQ files were quality controlled, aligned and analysed for enrichment of TDP-43-sensitive genes and junctions as above.

#### Dataset 2

FASTQ files were processed using the RNA-seq nfcore pipeline (v3.10.1)^61,62^. The pipeline was executed with Nextflow (v22.10.3)^36^, with alignment and quantification performed against the human genome GRCh38 and annotation release 107 from Ensembl (equivalent to Gencode v41). Sources of variation were assessed by performing principal component analysis on the gene level expression, which resulted in sex of the donor being included as a covariate.

Differential splicing, and enrichment analysis of TDP-43-sensitive genes and junctions was performed as above.

### Mouse *LRRK2* p.G2019S Model: Tissue Processing and Transcriptomic

#### Mouse Handling and Brain Tissue Collection

*LRRK2* p.G2019S mice have been described previously^63^ and were maintained on the same C57Bl/6J background for >10 generations. Animal studies were approved by the Institutional Animal Care and Use Committee at University of Florida. All mice were kept on a reverse cycle (light on from 8:30 pm to 8:30 am) and single-sex group-housed in enrichment cages after weaning at post-natal day 21, in accordance with NIH guidelines for care and use of animals. For all experiments, 3-6-month-old male animals were used. For tissue collection, mice were deeply anaesthetized via inhalation of isoflurane and then intracardially perfused with ice-cold phosphate buffer saline (PBS) prior to microdissection. All tissue was snap frozen in liquid nitrogen and stored at -80 °C.

#### RNA-seq Library Preparation and Sequencing

*LRRK2* p.G2019S KI mouse whole brain samples and their respective controls (30-50mg), were sent to Novogene Ltd. (Cambridge, UK), where RNA was extracted using the TRIzol extraction method. RNA quantitation, integrity and purity analysis was performed using the Agilent 5400 Fragment Analyzer System (Agilent Technologies).

cDNA libraries were prepared using a proprietary kit (Novogene UK) as detailed above for the *LRRK2* p.G2019S human cohort.

Libraries were sequenced using the NovaSeq X (Illumina) to obtain 150bp paired-end reads to a depth of ∼50 million reads respectively per sample.

### Generation of Mouse TDP-43-Sensitive Junctions and Genes

A mouse specific TDP-43 KD database of junctions and genes was made by downloading FASTQ files from GEO, as above for human experiments using sra_tools, ncbi_tools and entrez-direct packages. The following TDP-43 KD studies were downloaded: PRJNA141971^25^, PRJNA275618^26^, PRJNA290407^27^ (hippocampal samples only), PRJNA615680^24^ (somatodendritic samples only), PRJNA720642^23^ (neuronal samples only). Any technical replicate FASTQs were merged and run with the nextflow pipeline adapted for mouse experiments:

https://github.com/Jbrenton191/RNAseq_splicing_pipeline/tree/mouse-version.

The genome GRCm39 and gencode annotation M32 were used as references for alignment. For each experiment, covariate correction and LeafCutter was then run to produce a list of junctions and genes significantly spliced upon TDP-43 KD.

### *LRRK2* p.G2019S Primary Mouse Neuronal Culture Generation

Cortical neurons were isolated from pups of either sex at P0; brains were removed and placed on ice Earle’s Balanced Salt Solution (EBSS) supplemented with 50μM HEPES (GIBCO), 200μM Glucose (Gibco) and 1% Penicillin-streptomycin (Gibco). Cortical tissues from genotypes were pooled, dissociated in papain with DNase (Worthington Biochemical) and seeded at 150,000 cells/well in twenty-four well plates coated with poly-d-lysine (Sigma; P7280) or as 800,000 cells/well in 6-well coated plates, in Neurobasal-A medium (GIBCO; 0.5 mM α-glutamine and 2% B27). From DIV4, 10% of fresh media was added every 4–5 days and cultures were maintained in 5% CO2 at 37 °C.

### *LRRK2* p.G2019S Primary Mouse Neuronal Culture Immunocytochemistry

Cortical neurons grown to DIV17-21 were fixed in 4% PFA/4% sucrose in PBS for 12 min at RTP, permeabilized with methanol (3 min at −20°C) and blocked in 10% normal goat serum (NGS) in PBS for 1hr at RTP. Coverslips were incubated with primary antibodies overnight with agitation at 4°C in PBST plus 2% NGS, washed in blocking buffer (10% NGS+PBS) for 10min at RTP, then incubated with secondary antibody (in PBST + 2% NGS) for 1.5h at RTP. Following extensive washes with PBS at RTP, coverslips were mounted in Prolong Gold Antifade (Thermo Fisher Scientific). Imaging was conducted 24 h after mounting using a 60x oil objective on an Olympus FluoView FV-1000 confocal laser scanning microscope.

### Primary *LRRK2* p.G2019S Mouse Neuronal Culture Nuclear isolation and SDS-PAGE

Mouse cortical neurons (described above) underwent nuclear and cytoplasmic extraction using the NUC-PER kit (Thermo Fisher Scientific) per the manufacturer’s instructions. All buffers were supplemented with protease and phosphatase inhibitors (Thermo Fisher Scientific Cat # 78441) prior to use.

Total protein from isolated fractions was quantified using a Pierce BCA Protein Assay Kit (Thermo Fisher Scientific Cat # 23225) before being supplemented with NuPage LDS sample buffer 4x (#NP0008), boiled for 5 min at 95°C and run on a Bio-Rad Criterion TGX polyacrylamide gel at 150 V for 58 min. Proteins were transferred to nitrocellulose on a Bio-Rad Trans-Blot Turbo transfer system at 20 V for 7 min. The nitrocellulose was blocked for 1 h in 50% TBS (20 mM Tris, 0.5 M NaCl), 50% Intercept (TBS) Blocking Buffer (LI-COR; 927-66003) and incubated overnight with primary antibodies in 50% TBS-T (20 mM Tris, 0.5 M NaCl, 0.1% Tween 20), 50% Intercept (TBS) Blocking Buffer (LI-COR; 927-66003) at 4°C. Following 3 × 5 min washes with TBS-T, the nitrocellulose was incubated at RTP with secondary antibodies for 1 h, washed 3 × 5 min in TBS-T and scanned on a ChemiDoc MP imaging platform (Bio-Rad).

Secondary antibodies were used at 1:10,000 dilution: IRDyes 800CW Goat anti-Rabbit IgG (LiCor; #926–32,211) and 680RD Goat anti-Mouse IgG (LiCor; #926–68, 070).

### Antibodies for *LRRK2* p.G2019S mouse model experiments

The following antibodies were used throughout the study. For western blot: mouse monoclonal against *GAPDH* (Thermo Fisher Scientific, MA5-15738), mouse monoclonal against VPS35 (Abnova, (M02), clone 2D3), mouse monoclonal against Histone H3 (Abcam, ab1220), rabbit polyclonal against *LRRK2* (Abcam, MJFF2 c41-2; ab133474), rabbit polyclonal against TDP-43 (Proteintech, 10782-2-AP), rabbit polyclonal against FUS (Proteintech, 11570-1-AP). For ICC: rabbit polyclonal against TDP-43 (Proteintech, 10782-2-AP), rabbit polyclonal against FUS (Proteintech, 11570-1-AP) and chicken polyclonal against MAP2 (Abcam, ab5392).

### *LRRK2* p.G2019S Mouse Model Statistical Analysis

Immunocytochemical and immunoblotting experiments examining TDP-43 and FUS were analysed by two-sided unpaired T-tests using the rstatix package (v0.7.2) in R. Immunoblotting experiments are normalised by nuclear protein levels (ratio of TDP-43 or FUS to Histone H3 expression). Mean values ± S.E.M are presented, as well as individual data points.

### Proteomics of hESC-Derived Dopaminergic Neurons (mDN dataset 1)

#### Mass Spectrometry Data Generation and Analysis

Midbrain dopaminergic neurons from three independent control lines and two independent lines of *LRRK2* p.G2019S (6 replicates per clonal line), were directly lysed by adding 100µL of 2% SDS lysis buffer (2% SDS (wt/vol) in 100mM TEABC (Triethyl ammonium bicarbonate buffer) with Roche mini-Protease and Phos-STOP inhibitors). The samples were then boiled at 95°C for 5 minutes and subjected to sonication using a Bioruptor sonicator. Protein amounts were estimated using BCA assay and 40µg of protein amount was processed for quantitative proteomic analysis. Sample preparation was carried out using Strap assisted micro-column method and on-column tryptic digestion was performed using Lys-C+Trypsin mix and the resulting peptides were eluted and subjected to vacuum dryness using Speedvac concentrator. The detailed protocol can be found in: https://www.protocols.io/view/sample-preparation-protocol-for-total-proteomic-an-x54v9jyo1g3e/v1

250ng of tryptic peptide digest was used for proteomic analysis. The mass spectrometry data was acquired using Orbitrap Astral mass spectrometer in-line with Vanquish-Neo nano-liquid chromatography system using 30 samples per day method. Peptides were resolved on an analytical column (Aurora Ultimate™ 25cm×75 μm C18 UHPLC column #AUR4-25075C18). The full scan was acquired using Orbitrap mass analyser at 240K resolution in the mass range of 380-908 m/z. The MS2 data was acquired using narrow-window data independent acquisition 2m/z isolation widths. The raw mass spectrometry data was processed using the DIA-NN 1.8.1 software suite^64^ for database searches against the Human Uniprot database (02/2023). Library free search performed with default DIA-NN settings: trypsin as a protease, 2 missed cleavages, oxidation of Met as variable modification and carbamidomethylation of Cys as fixed modification. MS1 mass tolerance was set to 10ppm and MS2 at 20 ppm, retention time dependent cross-run normalisation was enabled, any LC high accuracy quantification and smart profiling library generation was enabled. Data was filtered at 1% FDR.

#### Gene-Level Quantification and Differential Analysis

Peptide counts were summated by gene to obtain gene-level counts. To normalize for library size, total protein counts across all genes were summed, divided by the median library size of samples. A library scaling factor was generated by dividing each sample’s library size by the median library size. This scaling factor was multiplied to all gene counts for each sample.

To discover genes with differential protein expression, t-tests were performed between control and *LRRK2* p.G2019S samples and FDR-corrected by the number of genes tested. Genes with an FDR-corrected p value below 0.05 were counted as differentially expressed. Genes were classified as significantly upregulated if their Benjamini and Hochberg FDR adjusted p-value was ≤ 0.05 and Log_2_ fold change was > 0, and significantly downregulated if their FDR ≤ 0.05 and Log_2_ fold change was < 0.

#### DESeq2 Differential Gene Expression

Differential gene expression analysis was performed with DESeq (v1.42.1) using replicate-collapsed gene count matrices and covariates generated as above. Non-protein coding genes, genes with 0 counts in at least one sample in either the *LRRK2* p.G2019S or control groups, as well as any gene overlapping ENCODE blacklist regions of the genome were removed prior to analysis. Genes were classified as significantly upregulated if their Benjamini and Hochberg FDR adjusted p-value was ≤ 0.05 and Log_2_ fold change was > 0, and significantly downregulated if their FDR ≤ 0.05 and Log_2_ fold change was < 0.

#### TDP-43-dependent mis-splicing effects on mRNA and protein expression

Genes that were differentially spliced between *LRRK2* p.G2019S and control mDNs, also mis-spliced upon TDP-43 loss of function, and showed differential expression at both the mRNA and protein levels were selected to assess the impact of mis-splicing. These candidate genes were grouped based on the direction of their mRNA and protein expression changes. A one-sided Fisher’s exact test was used to determine whether more genes than expected showed downregulation at both the mRNA and protein levels.

#### Prediction of Cryptic Peptides from Novel Splicing Events

Cryptic peptide sequences were predicted from junctions upregulated and differentially spliced in the LRRK2 p.G2019S vs control mDN1 analysis, as well as in TDP-43 KD experiments (mean dPSI > 0 across experiments). Only junctions that contained a novel donor or novel acceptor sequence as annotated by dasper (v1.5.3, junction_annot function; Gencode v41 reference annotation/GTF) were used to reliably identify transcripts from which to generate ORFs. Any transcript containing the annotated side of the junction was then converted to a nucleotide sequence with the novel position included using the Biostrings (v2.70.3), BSgenome.Hsapiens.UCSC.hg38 (v.1.4.5), factR (v1.4.0) and rtracklayer (v1.62.0) packages to generate sequences and import references. The ORFik package (v1.22.2) was then used to find the longest possible open reading frame of the novel junction-containing transcript. A CDS and subsequent amino acid sequence was then generated from this ORF for the novel transcript. Sequences for annotated transcripts, containing both annotated sides of junctions were generated alongside novel transcripts.

#### Detection of Cryptic Peptides in Mass Spectrometry Data

Predicted novel splice donor and acceptor sequences were used as search targets for mass spectrometry-derived peptides using DIA-NN^64^ (v1.8.1) in library-free mode with default settings. Peptide ion sequences not present in the Human UniProt database (February 2023 release) were aligned to novel and annotated transcripts to identify potential novel coding regions processed using in-house developed tool named “Cauldron” (https://github.com/noatgnu/cauldron). These aligned sequences were then re-aligned to novel and annotated transcripts using the pairwiseAlignment function from the Biostrings package. A normalized alignment score difference of ≥ 0.5 between novel and annotated transcripts was used as the threshold to classify peptides as mapping to novel insertions. Peptides were further filtered to include only those between 7 and 30 amino acids in length.

To determine if there was significant upregulation of these peptides in LRRK2 p.G2019S compared to controls,, half the minimum peptide ion count across all samples was added to each value, followed by normalization using sample-specific library size scaling factors. Mean log₂ fold changes between LRRK2 and control samples were calculated, and t-tests were performed to identify significantly upregulated cryptic peptides. P-values were FDR-corrected for multiple testing, and peptides were considered high-confidence if they had a log₂ fold change > 0 and an FDR-adjusted p-value < 0.05.

### Plot generation

Plots were made using ggplot (v3.5.1) and ggpubr (v0.6.0). Diagrams were made using Biorender. Images were edited in Inkscape (v1.3.1).

### Data and code availability

Raw FASTQ files for mid-stage and late-stage idiopathic datasets are available upon request through the ASAP CRN Cloud platform^65^: https://cloud.parkinsonsroadmap.org/collections.

## Acknowledgements

This research was funded in part by Aligning Science Across Parkinson’s through the Michael J. Fox Foundation for Parkinson’s Research (MJFF). We would like to thank UCL Genomics for the high quality sequencing of all post-mortem samples, and Novogene UK for the high quality sequencing of the *LRRK2* p.G2019S post-mortem and mouse samples. Urša Šušnjar and Emanuele Buratti, as well as James Shen and Liency Wu for their help and time to access and interrogate their datasets. The authors thank Shannon Wall and Alyssa Dellutri for their efforts toward primary neuron immunocytochemistry, microscopy and analyses, along with the animal care staff at the McKnight Brain Institute and the Cancer and Genetics Research Complex, University of Florida, Gainesville, Florida, USA.

---

**Extended Data Fig 1.**
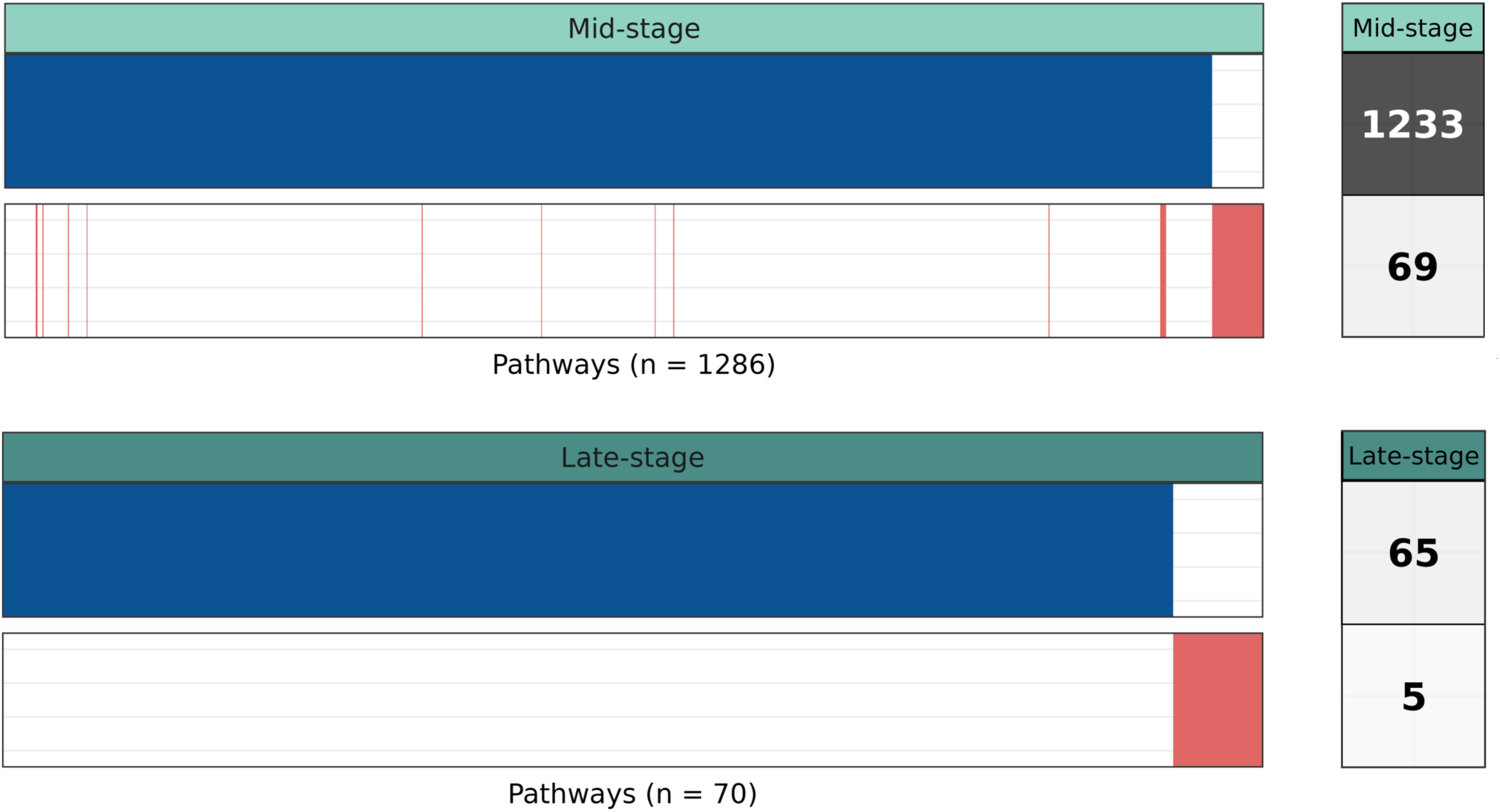
Overlap of significant pathways generated through differential expression or differential splicing analyses. Binary plot of common pathways found through DE and DS analyses in sporadic PD (FDR < 0.05). Each bar represents a single unique pathway and rows indicate differentially expression (blue) or splicing (red) at different pathological stages of PD (panels). Total numbers of unique pathways found for each stage and analysis shown to the right.

**Extended Data Fig. 2.**
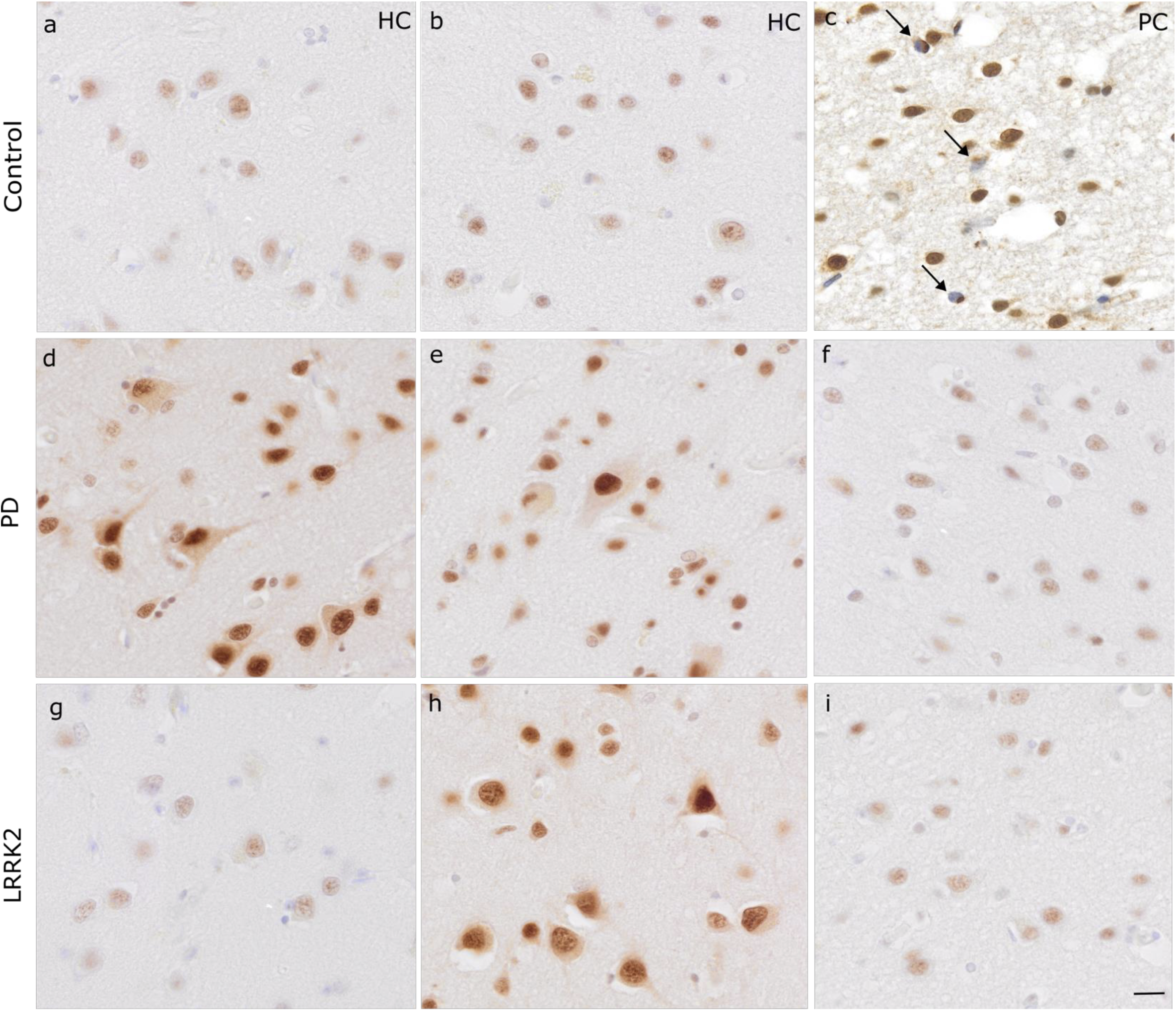
Absence of aggregated TDP43 in PD and LRRK2 cases. Representative images at 2Ox magnification highlighting TDP-43 staining (proteintech 10782-2-AP) of a-b, healthy control cases (HC); **c**, positive FTLD-TDP A case (PC); **d-f,** late-stage PD cases; **g-i,** *LRRK2* p.G2OI9S cases. Black arrows in panel d highlight aggregated TDP-43. Scale bar represents 2Oµm. Staining intensity varies with this antibody with varying fixation times of post-mortem cases.

**Extended Data Fig. 3.**
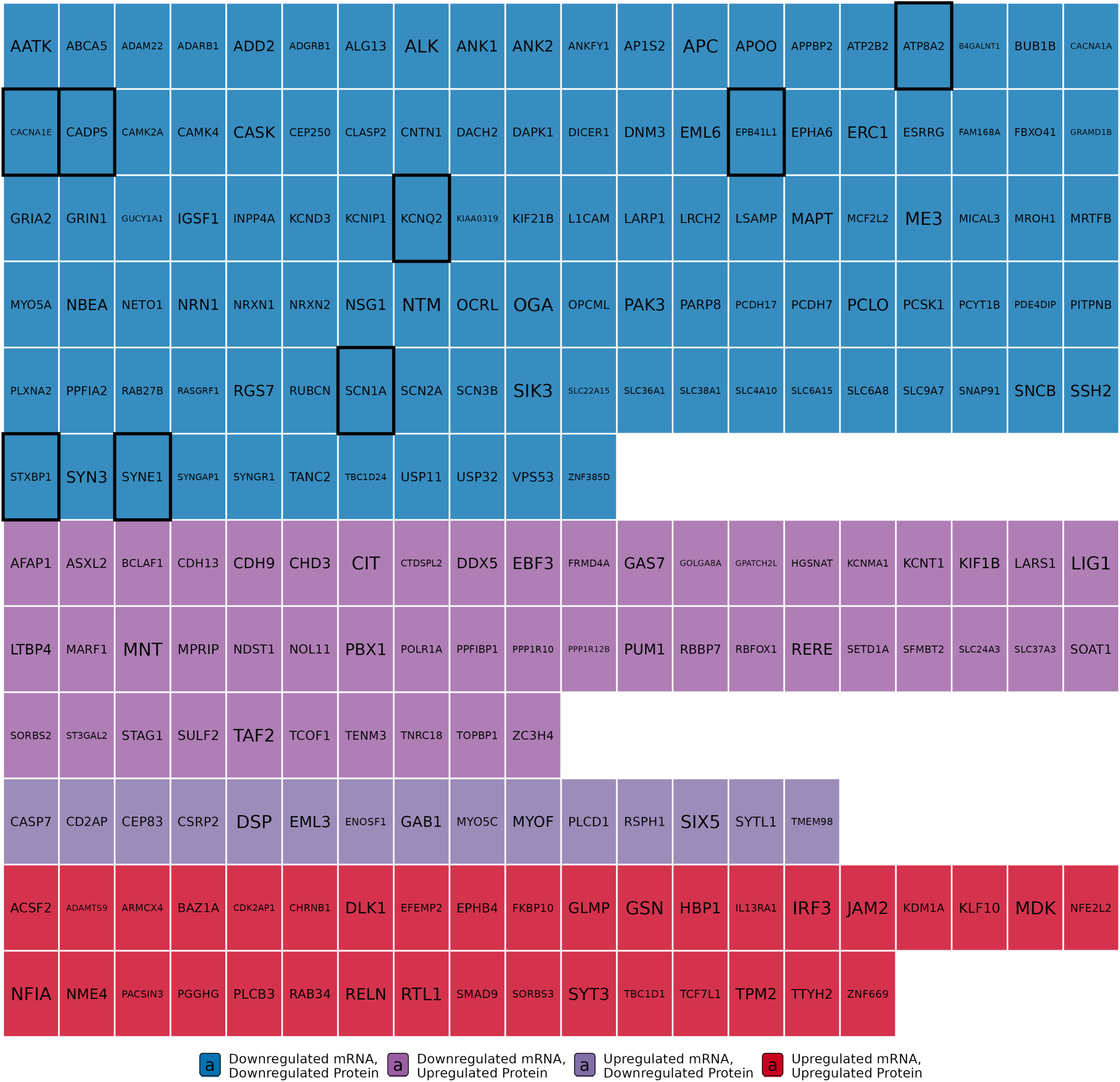
TDP-43-dependent mis-splicing leads to a predominant downregulation of protein and transcript expression in target genes. Waffle plot showing each gene that was a target of TDP-43 mis-splicing and had both significantly altered protein and gene expression as individual squares coloured by the direction of protein and transcript regulation. Genes outlined in black were previously identified as mis-spliced^28^, supporting the conclusion that TDP-43-mediated mis-splicing predominantly leads to downregulation at both the transcript and protein levels.

**Extended Data Fig. 4.**
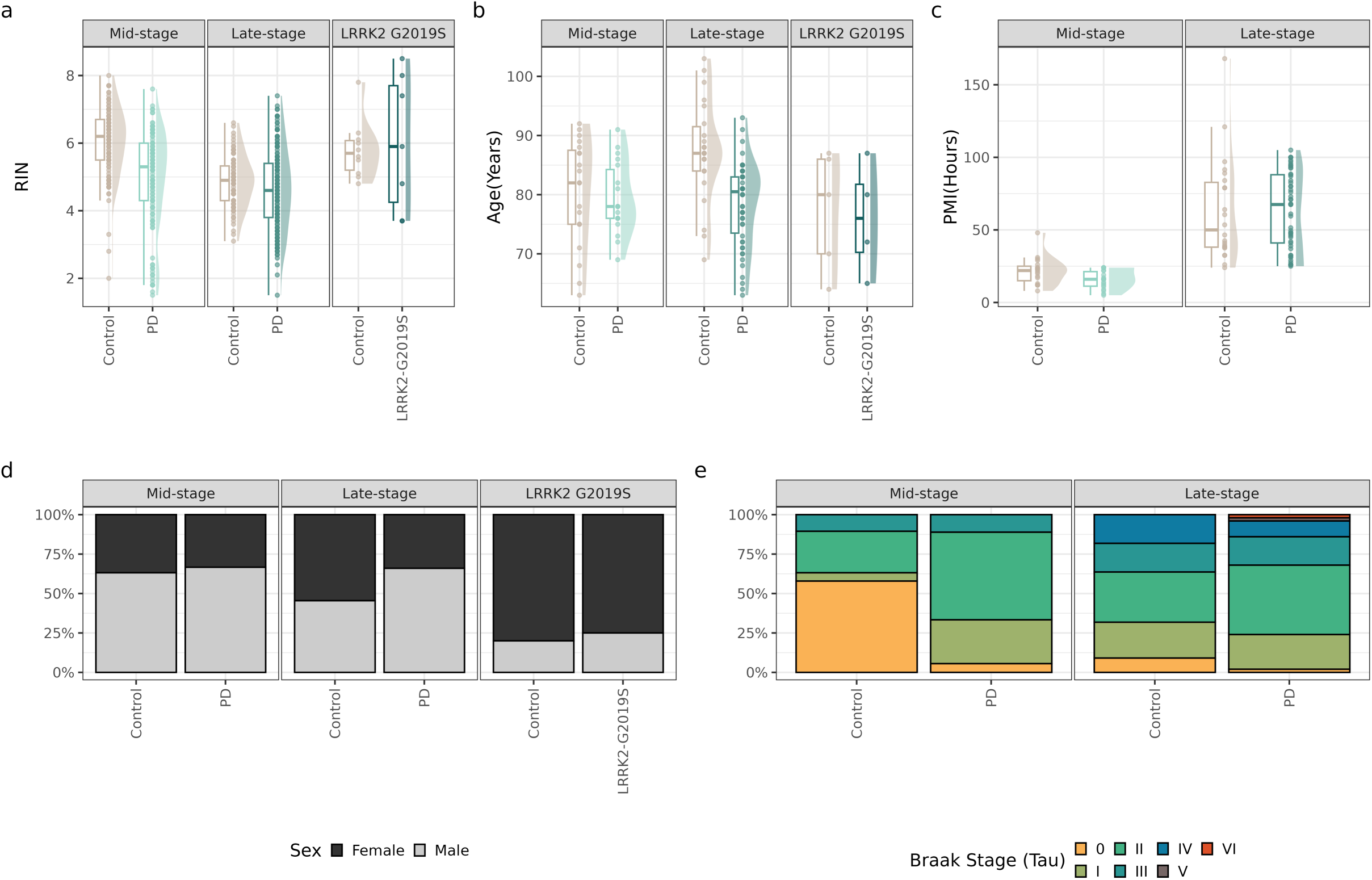
Demographics and metadata of post-mortem cohorts. Continuous sample and subject demographics: **a,** RNA integrity value (RIN); **b,** age of death of donor and **c,** post-mortem interval, and categorical demographics: **d.** sex and **e.** Braak neurofibrillary tangle staging, shown for mid-stage, late-stage and *LRRK2* p.G2OI 9S PD cohorts.

**Supplementary Table 1.** Sample numbers for each post-mortem cohort. Sample sizes for each brain region analysed in post-mortem datasets, with control cases listed first followed by Parkinson’s disease cases.

|  | Mid-stage | Late-stage | *LRRK2* p.G2019S |
| --- | --- | --- | --- |
| Substantia nigra | 9, 12 | NA | NA |
| Caudate | 15, 17 | NA | NA |
| Putamen | 13, 15 | NA | NA |
| Anterior cingulate cortex | 16, 16 | 21, 48 | 5, 3 |
| Temporal cortex | 15, 16 | 20, 47 | NA |
| Parahippocampal gyrus | 9, 15 | NA | NA |
| Frontal cortex | 18, 15 | 20, 48 | 5, 4 |
| Parietal cortex | 18, 15 | 19, 49 | NA |

**Supplementary Table 2.** Case demographics for TDP-43 immunohistochemistry. Age at death, age at onset and disease duration are given in years. PMI - Post Mortem Interval, this is given to the nearest hour. LB - Lewy body staging. SVD - Small vessel disease. AGD - Agyrophilic grain disease. AD - Alzheimer’s disease. CAA - Cerebral Amyloid Angiopathy.

| Case ID | Neuropathological diagnosis | Mutation | Age at onset | Age at death | Sex | Disease Duration | PMI | Braak LB | Braak Tau | CERAD | Thal | Confounding pathology | TDP-43 present | TDP-43 aggregation present |
| --- | --- | --- | --- | --- | --- | --- | --- | --- | --- | --- | --- | --- | --- | --- |
| 1 | Control | - | - | 81 | F | - | 32 | 0 | 3 | 0 | 0 | SVD | + | - |
| 2 | Control | - | - | 87 | F | - | 79 | 0 | 3 | 3 | 2 | Pathological aging, mild AGD | + | - |
| 3 | Control | - | - | 80 | F | - | 49 | 0 | 2 | 1 | 0 | Pathological aging, mild AGD, SVD | + | - |
| 4 | Control | - | - | 70 | F | - | 27 | 0 | 1 | 0 | 0 |  | - | - |
| 5 | Control | - | - | 86 | F | - | 40 | 0 | 0 | 0 | 0 | SVD | + | - |
| 6 | Parkinson's disease | - | 47 | 78 | F | 31 | 46 | 6 | 1 | 3 | 3 | Low level AD | ++ | - |
| 7 | Parkinson's disease | - | 41 | 63 | M | 22 | 38 | 6 | 1 | 3 | 3 | Low level AD | ++ | - |
| 8 | Parkinson's disease | - | 59 | 83 | F | 24 | 38 | 6 | 1 | 0 | 0 | PART | + | - |
| 9 | Parkinson's disease | - | 63 | 93 | F | 30 | 52 | 6 | 2 | 4 | 4 | Low level AD | + | - |
| 10 | Parkinson's disease | - |  |  |  |  |  |  |  |  |  |  |  |  |
| 11 | Parkinson's disease | G2019S LRRK2 |  | 84 | F |  | 32 |  |  |  |  |  | - | - |
| 12 | Parkinson's disease | G2019S LRRK2 | 48 | 87 | F | 39 | 43 |  | 2 |  | 1 | TDP-43 proteinopathy, low level AD, CAA | + | - |
| 13 | Parkinson's disease with dementia | G2019S LRRK2 | 51 | 80 | F | 29 | 44 |  |  |  |  |  | ++ | - |
| 14 | Parkinson's disease | G2019S LRRK2 | 57 | 72 | F | 15 | 25 |  | 2 |  |  | Low level AD | + | - |
| 15 | Parkinson's disease | G2019S LRRK2 | 54 | 65 | M | 11 |  |  |  |  |  |  | + | - |

**Supplementary Table 3.** Metadata for LRRK2 G2019S hiPSC-derived mDN dataset 2.

| Line | Induction | Mutation | Sex of donor | Genotype | Source | Age Of Donor | RRID |
| --- | --- | --- | --- | --- | --- | --- | --- |
| 41 | 1 | G2019S | Male | Mutant | NINDS | 52 | CVCL_QX27 |
| 346 | 1 | G2019S | Male | Mutant | UCL IoN iPS Bank | 68 | CVCL_WP53 |
| Ctrl4 | 1 | WT | Male | Control | Cedars Sinai SC02iCTR-NTn4 | 60-64 | NA |
| 41 | 2 | G2019S | Male | Mutant | NINDS | 52 | CVCL_QX27 |
| Ctrl1 | 2 | WT | Male | Control | In house (Tilo Lab) | 78 | CVCL_C6PM |
| Ctrl3 | 2 | WT | Female | Control | Thermo Fisher Scientific A18945 | Unknown | CVCL_RM92 |
| Ctrl4 | 2 | WT | Male | Control | Cedars Sinai SC02iCTR-NTn4 | 60-64 | NA |
| 41 | 3 | G2019S | Male | Mutant | NINDS | 52 | CVCL_QX27 |
| 346 | 3 | G2019S | Male | Mutant | UCL IoN iPS Bank | 68 | CVCL_WP53 |
| Ctrl1 | 3 | WT | Male | Control | In house (Tilo Lab) | 78 | CVCL_C6PM |
| Ctrl3 | 3 | WT | Female | Control | Thermo Fisher Scientific A18945 | Unknown | CVCL_RM92 |
| Ctrl4 | 3 | WT | Male | Control | Cedars Sinai SC02iCTR-NTn4 | 60-64 | NA |

## References

1. Buniello, A. et al. Open Targets Platform: facilitating therapeutic hypotheses building in drug discovery. Nucleic Acids Res. 53, D1467–D1475 (2025).

2. Ling, S.-C., Polymenidou, M. & Cleveland, D. W. Converging mechanisms in ALS and FTD: disrupted RNA and protein homeostasis. Neuron 79, 416–438 (2013).

3. Brown, A.-L. et al. TDP-43 loss and ALS-risk SNPs drive mis-splicing and depletion of UNC13A. Nature 603, 131–137 (2022).

4. Kapeli, K. et al. Distinct and shared functions of ALS-associated proteins TDP-43, FUS and TAF15 revealed by multisystem analyses. Nat. Commun. 7, 12143 (2016).

5. Klim, J. R. et al. ALS-implicated protein TDP-43 sustains levels of STMN2, a mediator of motor neuron growth and repair. Nat. Neurosci. 22, 167–179 (2019).

6. Melamed, Z. et al. Premature polyadenylation-mediated loss of stathmin-2 is a hallmark of TDP-43-dependent neurodegeneration. Nat. Neurosci. 22, 180–190 (2019).

7. Liu, E. Y. et al. Loss of Nuclear TDP-43 Is Associated with Decondensation of LINE Retrotransposons. Cell Rep. 27, 1409–1421.e6 (2019).

8. Roczniak-Ferguson, A. & Ferguson, S. M. Pleiotropic requirements for human TDP-43 in the regulation of cell and organelle homeostasis. Life Sci. Alliance 2, (2019).

9. Bandres-Ciga, S. et al. Large-scale pathway specific polygenic risk and transcriptomic community network analysis identifies novel functional pathways in Parkinson disease. Acta Neuropathol. 140, 341–358 (2020).

10. Keuss, M. J., et al. Loss of TDP-43 induces synaptic dysfunction that is rescued by UNC13A splice-switching ASOs. BioRxiv (2024) doi:10.1101/2024.06.20.599684.

11. Diaper, D. C. et al. Loss and gain of Drosophila TDP-43 impair synaptic efficacy and motor control leading to age-related neurodegeneration by loss-of-function phenotypes. Hum. Mol. Genet. 22, 1539–1557 (2013).

12. Agin-Liebes, J. et al. Patterns of TDP-43 Deposition in Brains with LRRK2 G2019S Mutations. Mov. Disord. 38, 1541–1545 (2023).

13. Wider, C., Dickson, D. W. & Wszolek, Z. K. Leucine-rich repeat kinase 2 gene-associated disease: redefining genotype-phenotype correlation. Neurodegener Dis 7, 175–179 (2010).

14. Garrido, A. et al. α-synuclein RT-QuIC in cerebrospinal fluid of LRRK2-linked Parkinson’s disease. Ann. Clin. Transl. Neurol. 6, 1024–1032 (2019).

15. Chahine, L. M. et al. LRRK2-associated parkinsonism with and without in vivo evidence of alpha-synuclein aggregates: longitudinal clinical and biomarker characterization. Brain Commun. 7, fcaf103 (2025).

16. Sun, M. et al. Cryptic exon incorporation occurs in Alzheimer’s brain lacking TDP-43 inclusion but exhibiting nuclear clearance of TDP-43. Acta Neuropathol. 133, 923–931 (2017).

17. Torres, P. et al. Selected cryptic exons accumulate in hippocampal cell nuclei in Alzheimer’s disease with and without associated TDP-43 proteinopathy. Brain 143, e20 (2020).

18. Schapansky, J. et al. Familial knockin mutation of LRRK2 causes lysosomal dysfunction and accumulation of endogenous insoluble α-synuclein in neurons. Neurobiol. Dis. 111, 26–35 (2018).

19. Beccano-Kelly, D. A. et al. Synaptic function is modulated by LRRK2 and glutamate release is increased in cortical neurons of G2019S LRRK2 knock-in mice. Front. Cell. Neurosci. 8, 301 (2014).

20. Volta, M. et al. Initial elevations in glutamate and dopamine neurotransmission decline with age, as does exploratory behavior, in LRRK2 G2019S knock-in mice. eLife 6, (2017).

21. MacIsaac, S. et al. Neuron-autonomous susceptibility to induced synuclein aggregation is exacerbated by endogenous Lrrk2 mutations and ameliorated by Lrrk2 genetic knock-out. Brain Commun. 2, fcz052 (2020).

22. Ling, J. P., Pletnikova, O., Troncoso, J. C. & Wong, P. C. TDP-43 repression of nonconserved cryptic exons is compromised in ALS-FTD. Science 349, 650–655 (2015).

23. Šušnjar, U. et al. Cell environment shapes TDP-43 function with implications in neuronal and muscle disease. *Commun*. Biol. 5, 314 (2022).

24. Briese, M. et al. Loss of Tdp-43 disrupts the axonal transcriptome of motoneurons accompanied by impaired axonal translation and mitochondria function. Acta Neuropathol. Commun. 8, 116 (2020).

25. Polymenidou, M. et al. Long pre-mRNA depletion and RNA missplicing contribute to neuronal vulnerability from loss of TDP-43. Nat. Neurosci. 14, 459–468 (2011).

26. Amlie-Wolf, A. et al. Transcriptomic Changes Due to Cytoplasmic TDP-43 Expression Reveal Dysregulation of Histone Transcripts and Nuclear Chromatin. PLoS ONE 10, e0141836 (2015).

27. Jeong, Y. H. et al. Tdp-43 cryptic exons are highly variable between cell types. Mol. Neurodegener. 12, 13 (2017).

28. Ma, X. R. et al. TDP-43 represses cryptic exon inclusion in the FTD-ALS gene UNC13A. Nature 603, 124–130 (2022).

29. Grima, N. et al. Multi-region brain transcriptomic analysis of amyotrophic lateral sclerosis reveals widespread RNA alterations and substantial cerebellum involvement. Mol. Neurodegener. 20, 40 (2025).

30. Irwin, K. E. et al. A fluid biomarker reveals loss of TDP-43 splicing repression in presymptomatic ALS-FTD. Nat. Med. 30, 382–393 (2024).

31. Albagli, E. A., Calliari, A., Gendron, T. F. & Zhang, Y.-J. HDGFL2 cryptic protein: a portal to detection and diagnosis in neurodegenerative disease. Mol. Neurodegener. 19, 79 (2024).

32. Calliari, A. et al. HDGFL2 cryptic proteins report presence of TDP-43 pathology in neurodegenerative diseases. Mol. Neurodegener. 19, 29 (2024).

33. Mehta, P. R., Brown, A.-L., Ward, M. E. & Fratta, P. The era of cryptic exons: implications for ALS-FTD. Mol. Neurodegener. 18, 16 (2023).

34. Braak, H. et al. Staging of brain pathology related to sporadic Parkinson’s disease. Neurobiol. Aging 24, 197–211 (2003).

35. Feleke, R. et al. Cross-platform transcriptional profiling identifies common and distinct molecular pathologies in Lewy body diseases. Acta Neuropathol. 142, 449–474 (2021).

36. Di Tommaso, P. et al. Nextflow enables reproducible computational workflows. Nat. Biotechnol. 35, 316–319 (2017).

37. Frankish, A. et al. GENCODE: reference annotation for the human and mouse genomes in 2023. Nucleic Acids Res. 51, D942–D949 (2023).

38. Chen, S., Zhou, Y., Chen, Y. & Gu, J. fastp: an ultra-fast all-in-one FASTQ preprocessor. Bioinformatics 34, i884–i890 (2018).

39. Patro, R., Duggal, G., Love, M. I., Irizarry, R. A. & Kingsford, C. Salmon provides fast and bias-aware quantification of transcript expression. Nat. Methods 14, 417–419 (2017).

40. Andrews, S. FastQC: A Quality Control tool for High Throughput Sequence Data. (Babraham Bioinformatics, 2010).

41. Wang, L., Wang, S. & Li, W. RSeQC: quality control of RNA-seq experiments. Bioinformatics 28, 2184–2185 (2012).

42. Okonechnikov, K., Conesa, A. & García-Alcalde, F. Qualimap 2: advanced multi-sample quality control for high-throughput sequencing data. Bioinformatics 32, 292–294 (2016).

43. Broad Institute. Picard2019toolkit. (Broad Institute, 2019).

44. Ewels, P., Magnusson, M., Lundin, S. & Käller, M. MultiQC: summarize analysis results for multiple tools and samples in a single report. Bioinformatics 32, 3047–3048 (2016).

45. Brenton, J. Jbrenton191/RNAseq_splicing_pipeline: Human RNAseq Alignment and QC for Differential Expression and Splicing Analysis. Zenodo (2024) doi:10.5281/zenodo.13379191.

46. Ritchie, M. E., et al. limma powers differential expression analyses for RNA-sequencing and microarray studies. Nucleic Acids Res. 43, e47 (2015).

47. Langfelder, P. & Horvath, S. WGCNA: an R package for weighted correlation network analysis. BMC Bioinformatics 9, 559 (2008).

48. Amemiya, H. M., Kundaje, A. & Boyle, A. P. The ENCODE blacklist: identification of problematic regions of the genome. Sci. Rep. 9, 9354 (2019).

49. Law, C. W., Chen, Y., Shi, W. & Smyth, G. K. voom: Precision weights unlock linear model analysis tools for RNA-seq read counts. Genome Biol. 15, R29 (2014).

50. Li, Y. I., et al. Annotation-free quantification of RNA splicing using LeafCutter. Nat. Genet. 50, 151–158 (2018).

51. Gene Ontology Consortium et al. The Gene Ontology knowledgebase in 2023. Genetics 224, iyad031 (2023).

52. Yu, G., et al. GOSemSim: an R package for measuring semantic similarity among GO terms and gene products. Bioinformatics 26, 976–978 (2010).

53. Sayols, S. rrvgo: a Bioconductor package for interpreting lists of Gene Ontology terms. MicroPubl. Biol. 2023, (2023).

54. Koriath, C. A. M., et al. The clinical, neuroanatomical, and neuropathologic phenotype of TBK1-associated frontotemporal dementia: A longitudinal case report. Alzheimers Dement (Amst) 6, 75–81 (2017).

55. Rohrer, J. D., et al. Clinical and neuroanatomical signatures of tissue pathology in frontotemporal lobar degeneration. Brain 134, 2565–2581 (2011).

56. Busquets, O., et al. iSCORE-PD: an isogenic stem cell collection to research Parkinson’s Disease. BioRxiv (2025) doi:10.1101/2024.02.12.579917.

57. Li, H., et al. Highly efficient generation of isogenic pluripotent stem cell models using prime editing. eLife 11, (2022).

58. Mohieddin Syed, K., Busquets, O., Soldner, F. & Hockemeyer, D. Dopaminergic neuron differentiation v1. (2024) doi:10.17504/protocols.io.3byl4q8yovo5/v1.

59. Li, H., Busquets, O., Poser, S., Hockemeyer, D. & Soldner, F. Nucleofection (Amaxa) and electroporation (Biorad) of hPSCs. (2023).

60. Virdi, G. S., et al. Protein aggregation and calcium dysregulation are hallmarks of familial Parkinson’s disease in midbrain dopaminergic neurons. npj Parkinsons Disease 8, 162 (2022).

61. Patel, H., et al. nf-core/rnaseq: nf-core/rnaseq v3.10.1 - Plastered Rhodium Rudolph. Zenodo (2023) doi:10.5281/zenodo.7505987.

62. Ewels, P. A., et al. The nf-core framework for community-curated bioinformatics pipelines. Nat. Biotechnol. 38, 276–278 (2020).

63. Yue, M., et al. Progressive dopaminergic alterations and mitochondrial abnormalities in LRRK2 G2019S knock-in mice. Neurobiol. Dis. 78, 172–195 (2015).

64. Demichev, V., Messner, C. B., Vernardis, S. I., Lilley, K. S. & Ralser, M. DIA-NN: neural networks and interference correction enable deep proteome coverage in high throughput. Nat. Methods 17, 41–44 (2020).

65. Hardy, J., Wood, N., Ryten, M., Lee, M. & Bras, J. ASAP CRN Cloud PMDBS bulk RNAseq Collection. Zenodo (2024) doi:10.5281/zenodo.14373344.

